# The heterogeneity and effect of diabetes and hypertension on spectral characteristics of vasomotion

**DOI:** 10.1101/2023.07.10.548472

**Authors:** Liangjing Zhao, Shuhong Liu, Yang Liu, Hui Tang

## Abstract

Vasomotion refers to the spontaneous oscillation of blood vessels within a frequency range of 0.01 to 1.6 Hz. Various disease states, including hypertension and diabetes, have been associated with alterations in vasomotion at the finger, indicating potential impairment of skin microcirculation. Due to the non-linear nature of human vasculature, the modification of vasomotion may vary across different locations for different diseases. In this study, Laser Doppler Flowmetry was used to measure blood flow motion at acupoints LU8, LU5, SP6, and PC3 among 49 participants with or without diabetes and/or hypertension. Fast Fourier Transformation was used to analyze noise type while Hilbert-Huang Transformation and wavelet analysis were applied to assess Signal Noise Ratio (SNR) results. Statistical analysis revealed that different acupoints exhibit distinct spectral characteristics of vasomotion not only among healthy individuals but also among patients with diabetes and hypertension. The results showed strong heterogeneity of vasomotion among blood vessels, indicating that the vasomotion measured at a certain point may not reflect the real status of microcirculation.

## Introduction

Vasomotion refers to the change in the diameter of blood vessels, particularly arterial diameter ^1, 2^. It manifests as a rhythmic oscillation in blood vessels^3^. The earliest recorded use of a microscope dates back to 1622^4^, and following its invention, the first systematic study of the vascular system was conducted on the web of a frog’s foot^5^. Early microscopic studies revealed the presence of vasomotion in human blood vessels^6^, dog omentum^7^, and bat wings^8^.

For humans, arterial oscillations are influenced by a range of factors including heartbeat, respiration, intrinsic myogenic activity of vascular smooth muscle, sympathetic nerve activity, and activity of vascular endothelium^9–11^. These five activities result in vasomotion across five frequency bands (around 1.0, 0.3, 0.1, 0.03 and 0.01Hz) respectively^12^. Heartbeat and respiration influence the frequency bands around 1.0 and 0.3Hz of vasomotion^13^, while various mechanical stresses exerted on smooth muscle cells and endothelial cells also impact vasomotion^14^. Blood flow in vessels induces variations in pressure and changes in shear stress. Arterial contraction results in an increase in calcium (Ca^2+^) concentration in the smooth muscle cytosolic^15^. An increase in intraluminal pressure leads to an increase in Ca^2+^ concentration, causing vessel contraction, a phenomenon known as myogenic response^15–17^, it corresponds to the 0.1Hz frequency band. The sarcoplasmic reticulum intermittently releases Ca^2+^ to smooth muscle cells, activating a chloride channel in the plasma membrane^18^. This process causes cell depolarization, which opens Ca^2+^ channels in both active and quiescent cells, activating the sarcoplasmic reticulum in quiescent cells^19^. In terms of neurohumoral influence, Traube-Hering-Mayer waves exhibit oscillations of neural firing with periodic rhythms of around 0.1Hz, in addition to their influence on the 0.03Hz frequency domain. Mayer waves are transient oscillatory responses to hemodynamic perturbations^20, 21^. In endothelial cells, the endothelium-nitric oxide(NO)-cyclic guanosine monophosphate axis significantly influences vasomotion^22^. Some studies have shown that NO can inhibit vasomotion in certain arteries^23, 24^, while others have demonstrated that NO can enhance vasomotion in other vessels^25, 26^. Nitric oxide (NO) can regulate vasomotion in two ways. It may influence endothelial cells and affect vasomotion. Considering these factors, the vasomotor system is nonlinear. Besides, various measures, such as local thermal changes^27, 28^, ischemia^29^, or acidosis^30^, can induce abnormal vasomotor activity. These are the inherent factors that influence vascular movement. Additionally, many diseases contribute to microvascular dysfunction.

Diabetes is a disease characterized by abnormally elevated blood glucose levels and is associated with insulin deficiency^31^. Insulin plays a role in capillary recruitment^32^. A study by Michiel et al. showed that the effect of insulin on capillary recruitment is partly mediated by neurogenic vasomotor activity^33^. Another group suggested that vascular smooth muscle also plays a role in insulin-induced capillary recruitment^34^. In 1995, Stansberry et al. demonstrated that the vasomotor amplitude of the fingers of diabetic patients was significantly reduced compared to healthy individuals^35^.

Reduced vasomotor amplitude has been observed in both insulin-independent and insulin-dependent diabetes, a phenomenon that is exacerbated by the presence of neuropathy^36, 37^. Pathological vasomotion can occur even in the early stages of diabetes^38^. Metformin, a commonly used drug in the treatment of non-insulin-dependent diabetes, has been shown to delay pathological vasomotion in diabetes mellitus^39^. Capillary blood flow may increase in the early stages of diabetes but decrease in long-term diabetes^40^. Metformin can improve post-ischemic perfusion of the microvascular bed in diabetic patients. Microvascular dysfunction in diabetes can manifest as increased capillary pressure and permeability^40^, and metformin has demonstrated potential efficacy in reducing capillary permeability in diabetes^41^. Additionally, metformin can reduce insulin resistance, hyperinsulinemia, and obesity in diabetic patients^42^. In diabetic patients, abnormal intracellular Ca^2+^ metabolism has been observed in arteries ^43–46^. As previously discussed, Ca^2+^ concentration influences vasomotion, and thus abnormal Ca^2+^ concentration may result in abnormal vasomotion. Several studies have provided evidence that diabetes is often accompanied by elevated blood calcium levels^47–49^.

High blood pressure is widely recognized to cause damage to microvascular function and structure^50^. However, research also suggests that changes in the microvasculature may occur before hypertension is clinically diagnosed^51^. Peripheral resistance is distributed throughout arterioles, capillaries, and venules. A low capillary density can result in high blood pressure. Studies on the relationship between capillary density and arterial pressure suggest that microvascular dysfunction may develop before hypertension^52^. Evidence supports that microvascular dysfunction, including blood vessel thinning and arterial narrowing, contributes to high blood pressure. Antihypertensive therapy has been shown to have a positive effect on skin microcirculation in patients^50^. Some studies suggest that a lack of Ca^2+^ may play a role in the development of hypertension^53, 54^.

In summary, abnormal concentrations of calcium (Ca2+) may occur in patients with diabetes and hypertension. The concentration of Ca2+ primarily affects the intrinsic myogenic activity of vascular smooth muscle. There are no apparent abnormalities in heartbeat and breathing activities in patients with diabetes and hypertension. Therefore, this study focuses mainly on the frequency band around and below 0.1Hz. Vasomotor abnormalities are also present in many diseases, not just diabetes and hypertension. Obesity and its related metabolic consequences can result in microvascular dysfunction^55, 56^. Adiponectin has been identified as the link between obesity and microvascular vasomotion^57^. In some coronary artery diseases, vasomotor abnormalities are consistently present; for example, increases in coronary blood flow induced by acetylcholine^58^. Patients with hypercholesterolemia in systemic sclerosis develop microvascular dysfunction, which can be improved with simvastatin therapy^59^. Even in Alzheimer’s disease and cerebral amyloid angiopathy, there is evidence of impaired vascular reactivity^60^. The scope of this study was to focus on the effects of diabetes and hypertension on vasomotion. Therefore, in this study, subjects must be screened to minimize the impact of other diseases on the results. Noise plays a fundamental role in the biomedical systems, for example, the heartbeat of human could be modeled as pink noise^61^. Since the vasomotor system is non-linear, there is also noise in vasomotor system. Stochastic resonance (SR) is a phenomenon in which an optimal amount of added noise results in maximum signal enhancement, while further increases in noise intensity degrade signal quality^62^. The Signal to Noise Ratio (SNR) is the most common way to quantify stochastic resonance. As shown in the SNR vs. Noise chart, there is a sharp rise first, and then a slow decline after reaching the highest point, this is the hallmark of stochastic resonance. SR could enhance the transmission and detection of specific signals with an optimal level of noise input^63^. When random noise is added to a dynamic system, its sensitivity to weak signals increases. On the SNR versus noise plot, SR occurs when there is a rapid peak followed by a gentle decline, as shown in Figure 4A. In nature, SR is widely used by wildlife; for example, crayfish use it to detect water movement signals^62^, and the cricket cercal sensory system detects air disturbances^64^. Research has shown that stochastic mechanosensory stimulation can enhance the stability of preterm infants’ breathing^65^, and specific white noise signals can improve postural control in people with stroke or peripheral neuropathy^66^. Evidence suggests that in the human brain, neural activity uses SR to integrate signals from both eyes^67, 68^. In healthy adults, heart rate expressed in non-linear dynamics is normal^69–71^, but in abnormal situations, non-linear dynamics in the heart rate system are obscured^72, 73^, and SR may not occur. Although SR is used in medicine to treat some diseases, no studies have shown whether SR has a role in the vasomotor system. In this study, we will observe the emergence of SR for all subject groups. Since SR can only be observed by analyzing the trend of SNR versus noise, its occurrence can only be determined through artificial judgment. In this study, SNR is calculated and discussed to determine the impact of diabetes and hypertension on flow motion at different measurement locations with different subject groups.

## Materials and methods

The investigation was conducted in accordance with the Declaration of Helsinki, approved by the local ethics committee, and written informed consent was provided to the subjects.

### Design of the Study and Subject Selection

The aim of this study is to investigate the impact of diabetes and hypertension on vasomotion at different measurement locations. Some studies have shown that stimulating acupoints and non-acupoints produce different results. For example, manual acupuncture on a contralateral acupoint elicits a response, while the same stimulation on non-acupoints does not^74^. Needling on acupoints has a therapeutic effect on abdominal obesity compared to needling on non-acupoints^75^. Research on acupoints driving specific autonomic pathways has shown that different acupoints have different reactions to electroacupuncture stimulation^76^. Therefore, measurement locations were selected based on Traditional Chinese Medicine. The Sanyinjiao-SP6 acupoint can be used to treat diabetes and hypertension^77^. For acupoints on the arms, those on the same meridian vessel or those on different meridian vessels but in close proximity can be selected. The selected measuring acupoints should also be easy to locate. Jingqu-LU8 is near the pulse point on the wrist and is easy to find and measure. Quze-PC3 is located in the middle of the cubital crease and is easy to locate. Chize-LU5 is near Quze-PC3 and both Jingqu-LU8 and Chize-LU5 are on the lung meridian of hand-taiyin. Sanyinjiao-SP6 is an acupoint commonly used to treat diabetes and hypertension. Therefore, Jingqu-LU8, Quze-PC3, Chize-LU5, and Sanyinjiao-SP6 were selected as measurement locations, as shown in Figure 1, where LU-8, PC-3, S-P6, and LU-5 are acupoints according to the Standard of Acupuncture Nomenclature^78^. This study uses Laser Doppler Flowmetry (LDF) to measure skin perfusion. The laser Doppler technique calculates the skin perfusion index by measuring the Doppler frequency shift^79^. The results provide quantification of the product of flux, mean erythrocyte velocity, and concentration^80^. LDF measures small-volume, high-sampling blood flow by placing a probe on the skin surface^81^. It is non-invasive, fast, and less expensive than other measurement methods. Therefore, LDF is suitable for obtaining constant perfusion measurements and dynamic measurement capabilities in this study. This experiment was conducted in accordance with the Declaration of Helsinki and was approved by the local ethics committee. All subjects received detailed information about the procedures and purpose of the research and signed a consent form before participating. They were asked to provide truthful information. To obtain more accurate results and avoid the effects of other diseases, subjects with coronary, peripheral or cerebral artery diseases, heart failure or renal failure were excluded. All participants provided information about their gender, age, smoking status, and body mass index (BMI), as shown in Table 1.

**Figure 1.**
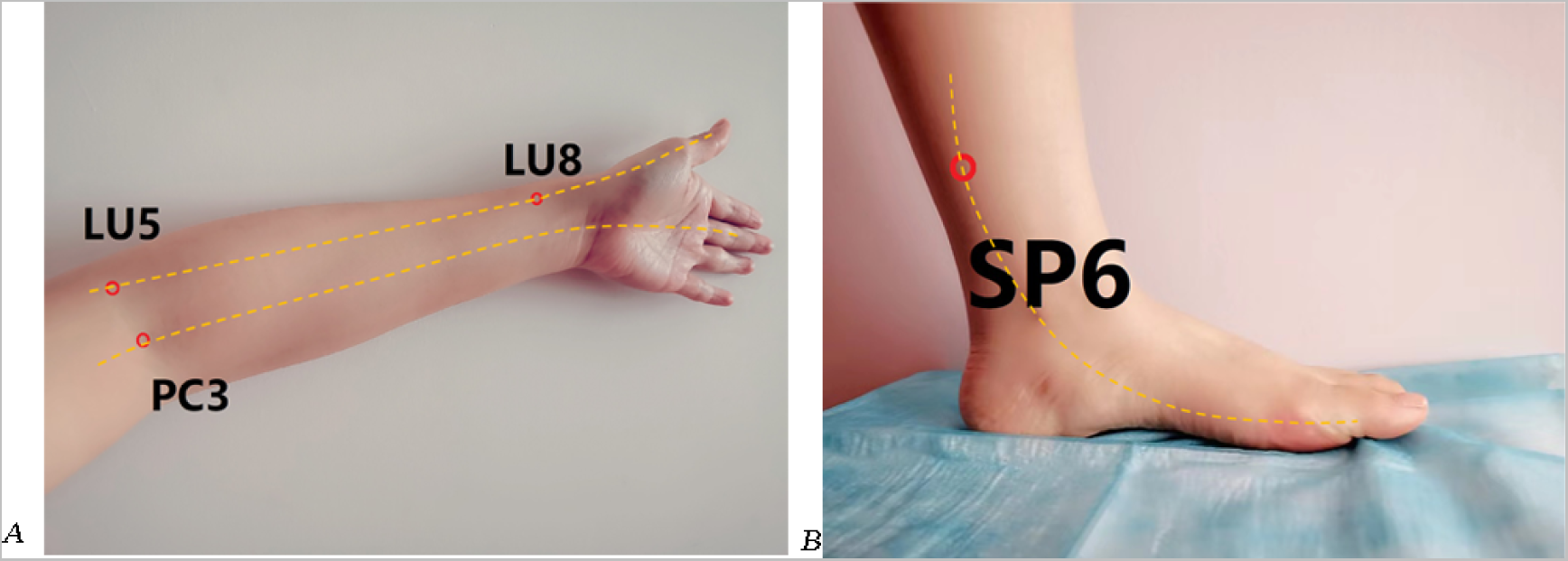
The Measurement Locations. (A) Jingqu-LU8, is at the depression between the radial styloid process and radial artery on the radial side of the palmar face of the forearm, which is on the lung meridian of hand-taiyin. Quze-PC3, is in the middle of the cubital crease, and is on the pericardium meridian of hand-jueyin. Chize-LU5, is at the radial depression of the biceps tendon and is on the lung meridian of hand-taiyin. (B) Sanyinjiao-SP6, is about four fingers upper of the medial malleolus tip and is at the posterior tibial edge near the bone edge depression. Sanyinjiao is on the spleen meridian of foot-taiyin.

**Table 1.** The Characteristics of the subjects

| Variable | Diabetes<br>(n=19)<br>Group D | Hypertension<br>(n=12)<br>Group H | Diabetes+<br>Hypertension<br>(n=13)<br>Group D+H | Control Group<br>(n=5)<br>Group C |
| --- | --- | --- | --- | --- |
| Female/male,<br>n(%) | 14(73.7)/5(26.3) | 7(72)/5(32) | 10(76.9)/3(23.1) | 5(100)/0(0) |
| Age, average<br>(extreme<br>values) years | 59(33-75) | 57.2(39-76) | 65(50-79) | 45.4(26-65) |
| Smoking, n(%) | 1(5.3) | 4(33.3) | 1(7.7) | 0(0) |
| BMI, average<br>(extreme<br>values) kg/m2 | 23.7(20.0-31.6) | 25.6(19.9-31.2) | 24.1(20.9-28.8) | 23.4(19.3-27.4) |

### Measurement Procedure

Before the measurement procedure, subjects were instructed to avoid consuming food, drugs, alcohol, coffee, and tea for at least 8 hours. They were required to lie in a supine position for at least 5 minutes before the measurement to acclimate to the environment. During the measurements, subjects were instructed to remain still and silent in a supine position. All measurements were conducted in the morning and took approximately 35 minutes per subject.

### Fast Fourier Transformation (FFT) & Noise type Analysis

The measured signal from LDF is time series, and FFT was used to Transform the time series signal into frequency domain^82^. FFT is widely used in describing the blood flow perfusion for different diseases, such as hypertension^83^, pulp necrosis^84^, retinopathy^85^ and so on.

Noise can be categorized into white noise and colored noise. White noise has equal frequency distribution over a wide range of frequencies with uniform intensity. Colored noise, on the other hand, has an uneven distribution of frequency components. Pink noise is a type of colored noise in which the intensity is inversely proportional to the frequency, resulting in equal energy across all octaves. In terms of power at a constant bandwidth, pink noise decreases at a rate of 3 dB per octave, and Brownian noise decreases at a rate of 6 dB per octave.

The color of noise can be determined from the FFT results by analyzing the slope of the fitting line, as shown in Figure 2B. The relationship between frequencies and octaves can be expressed using the definition of octaves:

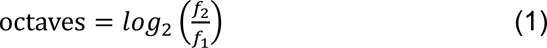

**Figure 2.**
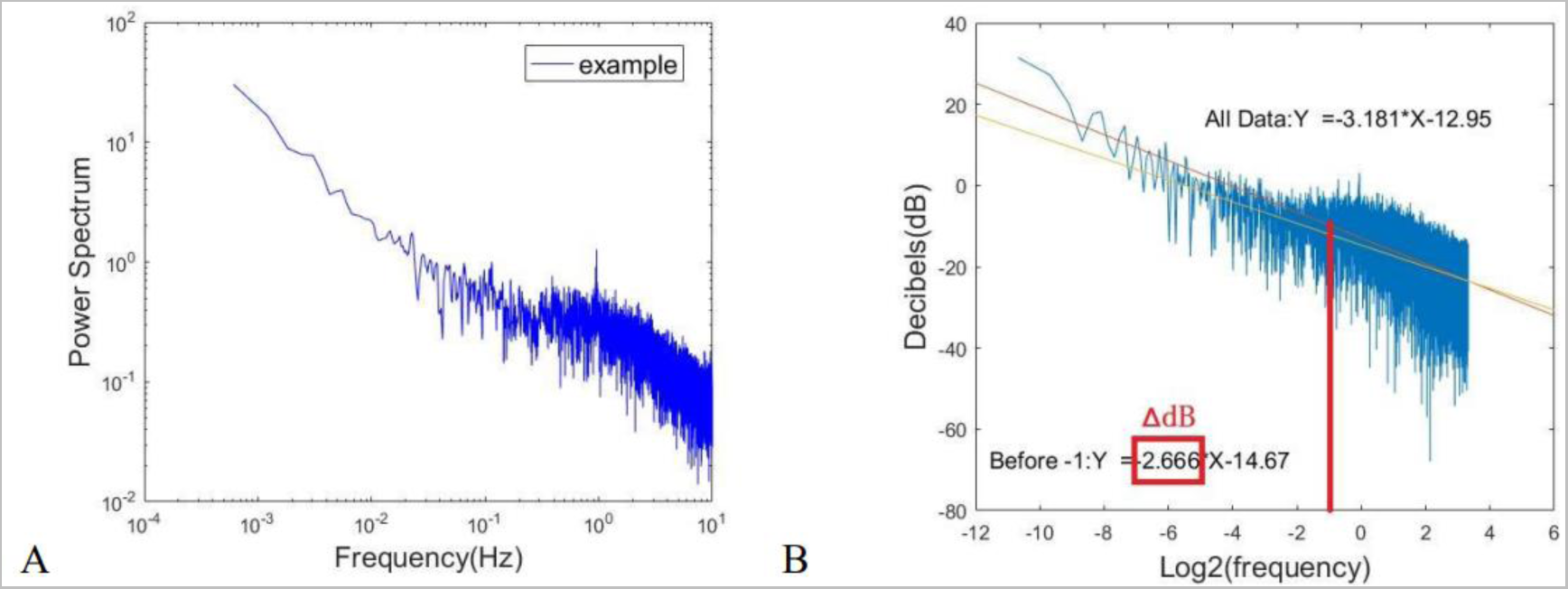
FFT Results of an example Time series. (A) The direct result from FFT for the example signal. (B) The results of noise type from FFT. There is a peak at around 1Hz from FFT, due to the activity of the heartbeat. To reduce the impact of this peak on the trend line, the noise type should use the trend line with x axis from minus infinity to −1. The red line is the trend line of the FFT results, and the orange line is the trend line of the results with x axis less than −1. 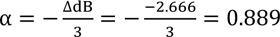, 0.889 is close to 1. Thus, for the example, the noise type is pink noise.

Assuming there is a noise, no matter if it is white, pink or Brownian noise, it should follow the equation:

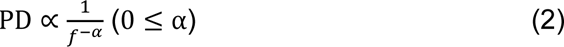

In this equation, α is the index criterion of the noise color. If α=0, the noise color is white, if α=1, the noise color is pink, and for α=2, the noise color is brown.

Where 𝑓_1_ and 𝑓_2_ are two different frequencies, 𝑃𝐷_2_ and 𝑃𝐷_1_ are the power spectral densities of 𝑓_1_ and 𝑓_2_.

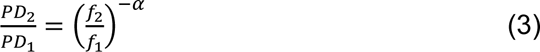

The definition of decibels gives the equation:

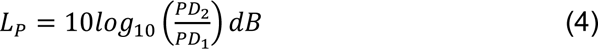

Where 𝐿_𝑃_ is the ratio of 𝑃𝐷_2_ and 𝑃𝐷_1_ expressed in dB.

It is known that when the pink noise is ideal, it means α = 1, and the power density drops at a rate of 3dB per octave.

When α = 1,

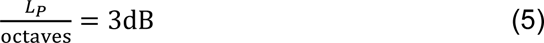

If the noise type is unknown,

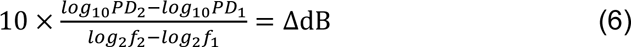

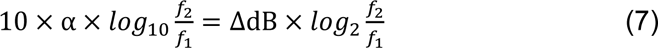

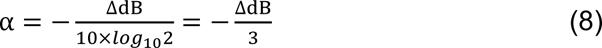

The value of α can be used as an indicator of the type of noise present. If α is around 2, it suggests that brown noise is dominant. If α is around 1, pink noise predominates in the frequencies. If α is around 0, the data represents white noise. Therefore, the type of noise can be determined by calculating the slope of the trend line on FFT results. There is often a peak near 1Hz on FFT results due to the influence of the heartbeat. This peak can significantly affect trendlines. Since our focus is on the low-frequency domain, we used trendlines for frequencies below 0.1Hz to calculate the type of noise. The results are based solely on the slope of the low-frequency domain.

### Wavelet, Hilbert-Huang Transformation (HHT) & SNR Analysis

FFT results only show the accumulated spectral power of a signal in the frequency domain. Wavelet transformation can reveal signal characteristics in both the time and frequency domains^86^. It is commonly used to analyze blood perfusion ^87–89^. Wavelet results are displayed in 3D, with spectral power represented by a color map, as shown in Figure 3A. In this study, the Morlet wavelet was chosen as the mother wavelet.

**Figure 3.**
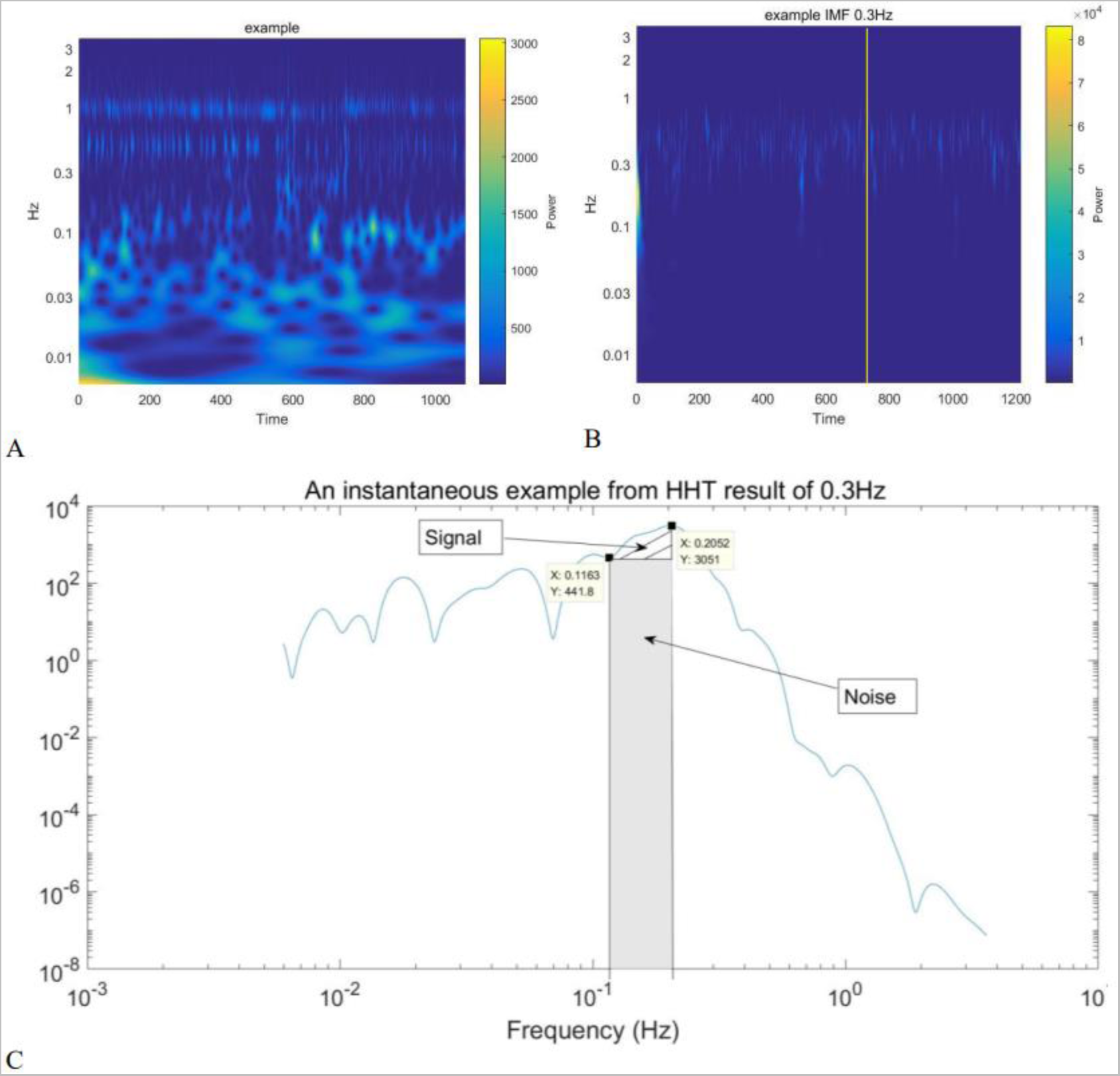
Example of HHT. (A) The wavelet result of the example signal. It shows all the frequency bands. (B) The HHT results of example for frequency bands 0.3Hz, and the instantaneous example selection. (C) An instantaneous example from HHT result of 0.3Hz. The peak point is the point at 0.2052Hz. The noise is the point with 0.1163Hz. 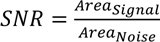.

**Figure 4.**
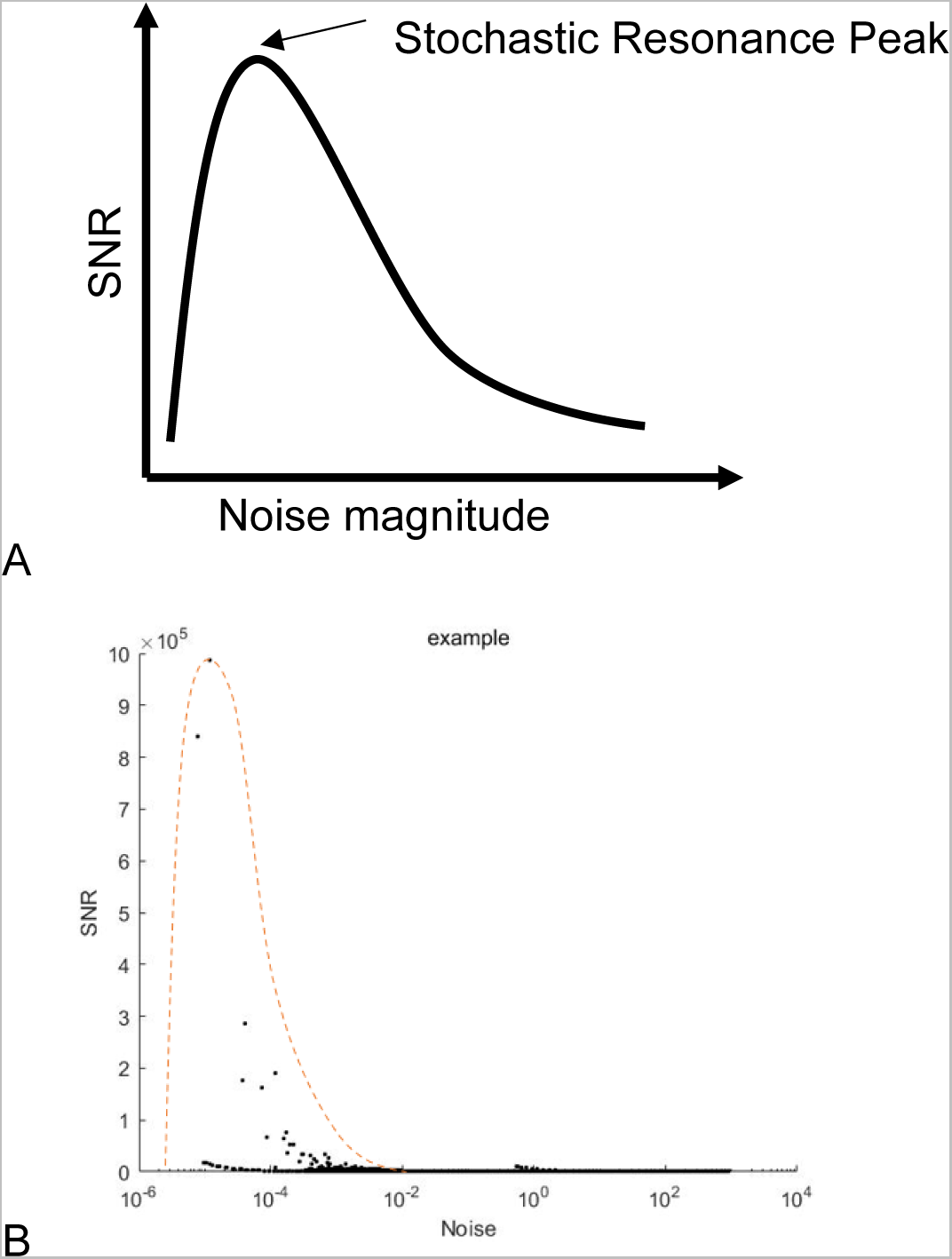
Example of SR. (A) The abridged general view of SR phenomenon. (B) The SNR to Noise chart of example signal. The SR phenomenon could be found in this chart.

Wavelet analysis produces a color map, as shown in Figure 3A. However, the boundaries between different frequency domains are not clearly defined on the color map of wavelet analysis results, particularly for frequencies below 0.1Hz. Additionally, different frequency domains can interfere with each other in instantaneous results. Therefore, HHT is used to decompose the original signal into signals in different frequency domains. Hilbert-Huang Transform consists of two main parts, the empirical mode decomposition and Hilbert spectral analysis^90–92^. The complex signals could be decomposed by empirical mode decomposition into several different frequencies signals, also could be called intrinsic mode functions (IMFs). HHT could give a better solution for the nonlinearity and nonstationary system than the traditional paradigm with constant frequency and amplitude.

In this experiment, IMFs are transformed using a wavelet to obtain spectral power in different domains. Only five frequency domains are studied in this experiment. For the frequency band around 1Hz, where the heartbeat is strong, the SNR is dominated by the intensity of the heartbeat, which may be influenced by cardiovascular health. At the same time, as mentioned before, the study mainly focuses on the frequency band around and below 0.1Hz, due to the possible abnormal concentration of Ca^2+^ in diabetes and hypertension. Therefore, SNR results for the frequency band around 1Hz were not discussed. However, the frequency domain of endothelial cell activity cannot be clearly distinguished from the frequency domain of sympathetic nerve activity using IMFs. As a result, the frequency domains of endothelial cells and sympathetic nerve activity were merged into a frequency band around 0.03Hz. To verify the presence of SR, frequency domains were manually selected. The instantaneous SNR was then extracted from the instantaneous Figure 3B showing the power spectral density versus frequency. After selecting the frequency domain (𝑓_𝑙𝑖_∼𝑓_𝑢𝑖_), the frequency of the peak point, 𝑓_𝑝𝑖_, is determined. The first concave vertex between 𝑓_𝑙𝑖_ and 𝑓_𝑝𝑖_ is then identified and denoted as (𝑓_𝑛𝑖_, 𝑃_𝑛𝑖_). The instantaneous SNR could be calculated as:

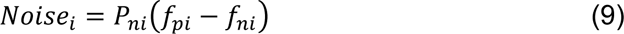

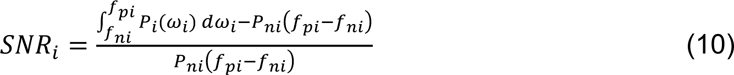

As shown in Figure 3C.

Plot all the instantaneous SNR versus instantaneous noise to verify the SR.

In this study, the SNR is calculated and analyzed to determine the impact of diabetes and hypertension on flow motion at various measurement locations across different subject groups. The results of the SNR analysis are then subjected to statistical evaluation.

### Statistical analysis

In the results section, data is presented as box plots for different groups and measurement locations. The top and bottom lines of the boxes represent the 75th and 25th percentiles, respectively, while the lines within the boxes represent the medians. The top and bottom lines outside the boxes represent the highest and lowest values, excluding outliers. Result tables display p-values. Data from different groups and measurement locations are compared in pairs using an unpaired t-test for statistical analysis. The confidence interval is 95%, it is means that the p-value of less than 0.05 was considered statistically significant in this study^93^. Both equal variance and unequal variance t-tests were used in calculations due to variance differences between groups.

## Results and discussion

### Results of Noise type

Figure 5 displays the values of α for different groups and measurement locations. As previously mentioned, the noise type indicated by the number α can be used to distinguish between noise colors. Figure 5 demonstrates the heterogeneity of noise color at different locations for various diseases. For PC-3 and LU-5 in different groups, there are significant differences in median and variance between groups. In contrast, LU-8 and SP-6 have similar median and variance values between groups. In Group D (diabetes), the median of LU-5 is higher than other measurement locations, and its variance is notably smaller than other measurement locations.

**Figure 5.**
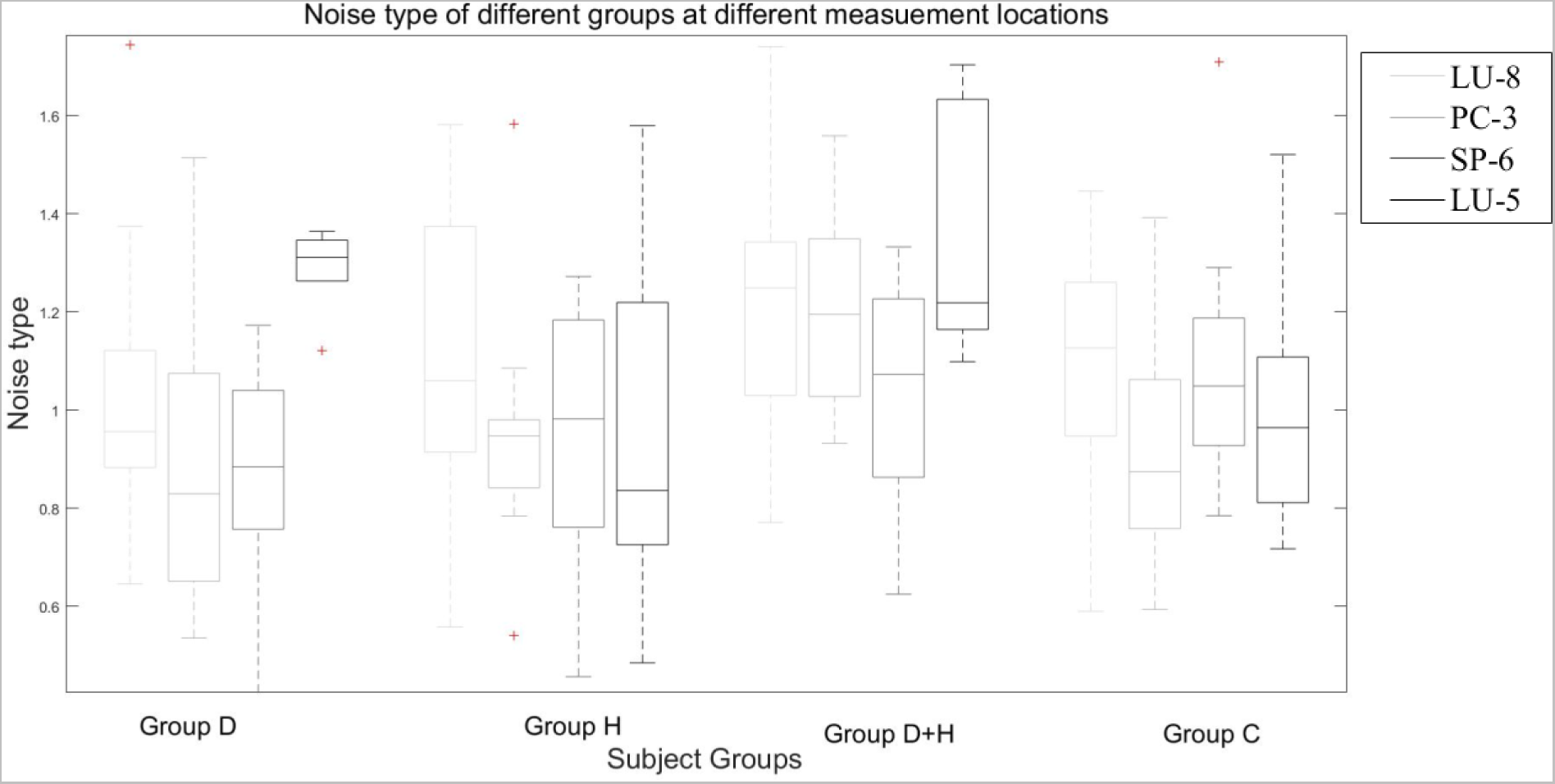
The results of Noise types of different groups at different measurement locations.

From Table 2, at location of LU-8, there is a significant difference between Group H (hypertension) and Group C (Control) at LU-8, where the *p* =.0182 < .05. For the Group H (hypertension) and Group C (control), *p* =.0209< .05. Even for the Group D+H (diabetes + hypertension) and Group C (control), *p* =.0000681< .05. These indicate that diabetes or hypertension has altered the noise color of flow motion signal significantly at LU-8.

**Table 2.** p-value of Noise type of different groups at different measurement locations.

| P-VALUE | LU-8 | PC-3 | SP-6 | LU-5 |
| --- | --- | --- | --- | --- |
| D/H | 0.181 | 0.0862 | 0.796 | 0.0575 |
| D/D+H | 0.0553 | **0.0125** | 0.0574 | 0.493 |
| D/C | **0.0182** | 0.303 | 0.273 | 0.407 |
| H/D+H | 0.838 | 0.637 | 0.0539 | 0.413 |
| H/C | **0.0209** | 0.941 | 0.407 | 0.668 |
| D+H/C | **0.000681** | 0.817 | **0.0189** | 0.859 |

At location of PC-3. The Group D+H (diabetes + hypertension) and Group D (diabetes), *p* = .0125< .05. And the Group H (hypertension) and Group D (diabetes), *p* = .0862 < .10. These forecast the hypertension may influence the noise color of flow motion signal significantly at PC-3.

At location of SP-6 there is a significant difference between Group D+H (diabetes + hypertension) and Group D (diabetes) at SP-6, where the *p* =.0574 < .10. For the Group D+H (diabetes + hypertension) and Group H (hypertension), *p* =.0539< .10. Even for the Group D+H (diabetes + hypertension) and Group C (control), *p* =.0189< .05. These indicate that the combination of diabetes and hypertension has altered the noise color of flow motion signal significantly at SP-6.

Therefore, diabetes would change the noise color at LU-8, the hypertension would change the noise color at LU-8 and PC-3, and hypertension +diabetes would change the noise color of flow motion at LU-8 and SP-6. This indicates that the noise color of flow motion signal at different location is different.

### Results of SR and SNR

The measured time series data is analyzed using HHT. One common method for quantifying stochastic resonance is through the SNR, which can be obtained from the output by forming the power spectrum. The power spectrum measures the frequency content of a time series^94^. Figure 6 displays several examples of how SNR varies with noise level. From Figure 6, SR can be found to occur in the vasomotor system for all the groups.

**Figure 6.**
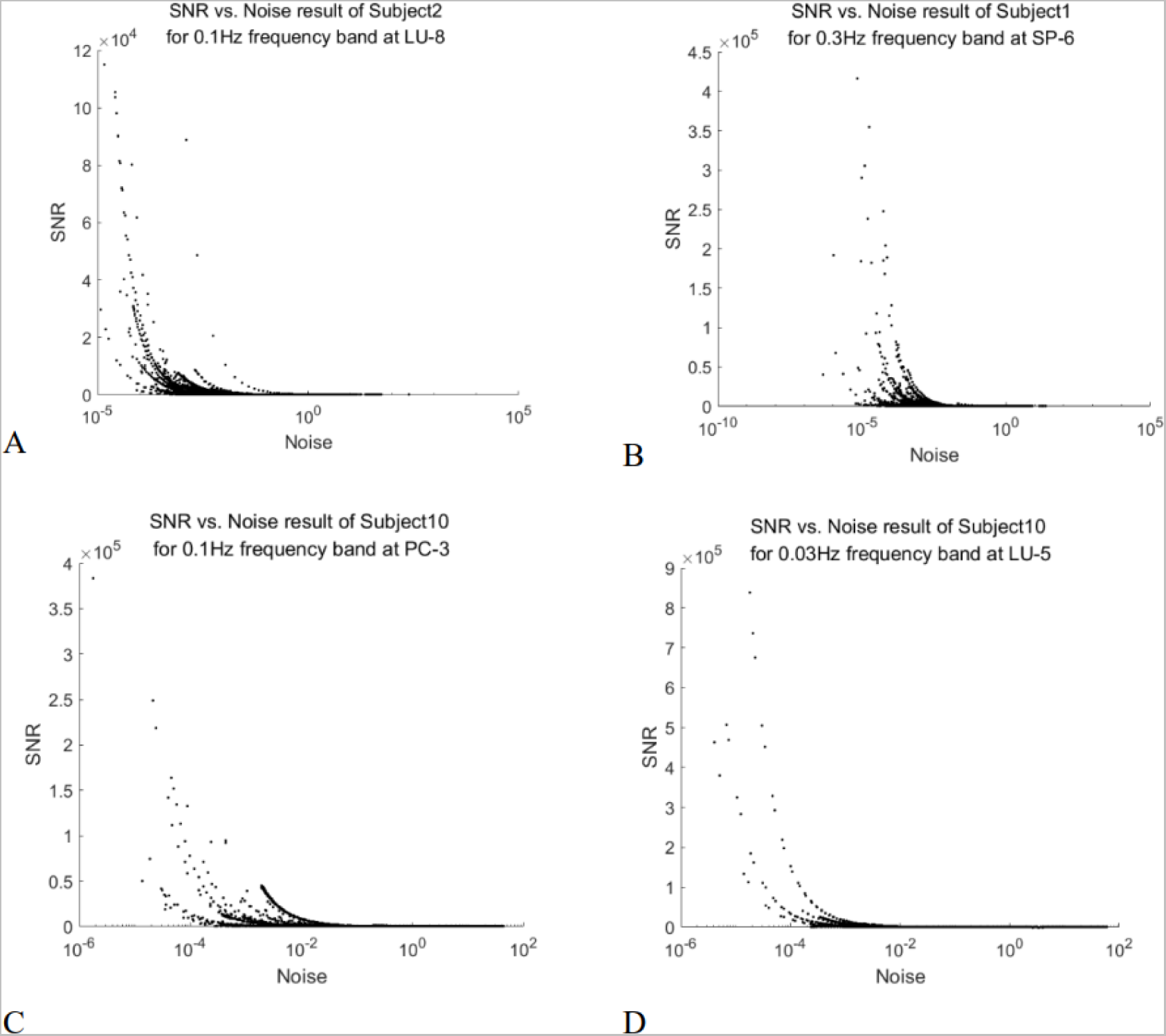
The example results of SNR vs. Noise figures. (A) The results of SNR vs. Noise figures for Subject 2 at 0.1Hz at LU-8. (B) The results of SNR vs. Noise figures for Subject 1 at 0.3Hz at SP-6. (C) The results of SNR vs. Noise figures for Subject 10 at 0.1Hz at PC-3. (D) The results of SNR vs. Noise figures for Subject 10 at 0.03Hz at LU-5.

The maximum value of the SNR can be used to describe certain features of the signal. However, in Figure 6C, the highest points appear to be singularities. To mitigate the potential influence of these singularities, the average value of the top 0.1%, 1%, 5%, and 10% of the SNR for each IMF is calculated to reduce their effects. The order of magnitudes between the top is large, so the convergence check is taken with the logarithmic function 𝑙𝑜𝑔_10_. The differences between the various top SNR could be calculated by:

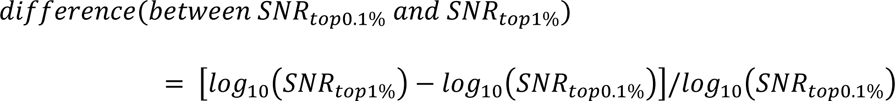

The rest can be done in the same manner.

The average top SNR results for four random IMFs were shown in the Table 3 below. The differences between the average SNR of the top 5% and 10% are less than 10%. It means that the effects of distorted points could be neglected for the average top 5% SNR.

**Table 3.** The average top SNR results for four random IMFs.

|  | **0.1%** | **1%** | **5%** | **10%** |
| --- | --- | --- | --- | --- |
| **Sample IMF1** | 79714.97 | 20108.44 | 5392.8 | 2865.229 |
| **Differences of Sample IMF1** | 12.2% | 13.3% | 7.4% | / |
| **Sample IMF2** | 57276.06 | 12455.97 | 3306.83 | 1749.811 |
| **Differences of Sample IMF2** | 14.0% | 14.1% | 7.9% | / |
| **Sample IMF3** | 60004.72 | 15926.54 | 4661.584 | 2512.743 |
| **Differences of Sample IMF3** | 12.1% | 12.7% | 7.3% | / |
| **Sample IMF4** | 811326.2 | 103785 | 23284.78 | 11936.58 |
| **Differences of Sample IMF4** | 15.1% | 12.9% | 6.6% | / |

Hence, the average SNR for the top 5% of full data for an IMF is used to discuss the discrepancy between different groups for all the measuring locations.

Figure 7 displays the SNR values for different groups and measurement locations at a frequency band of 0.3Hz. It illustrates the heterogeneity of SNR at different locations for various diseases at this frequency band.

**Figure 7.**
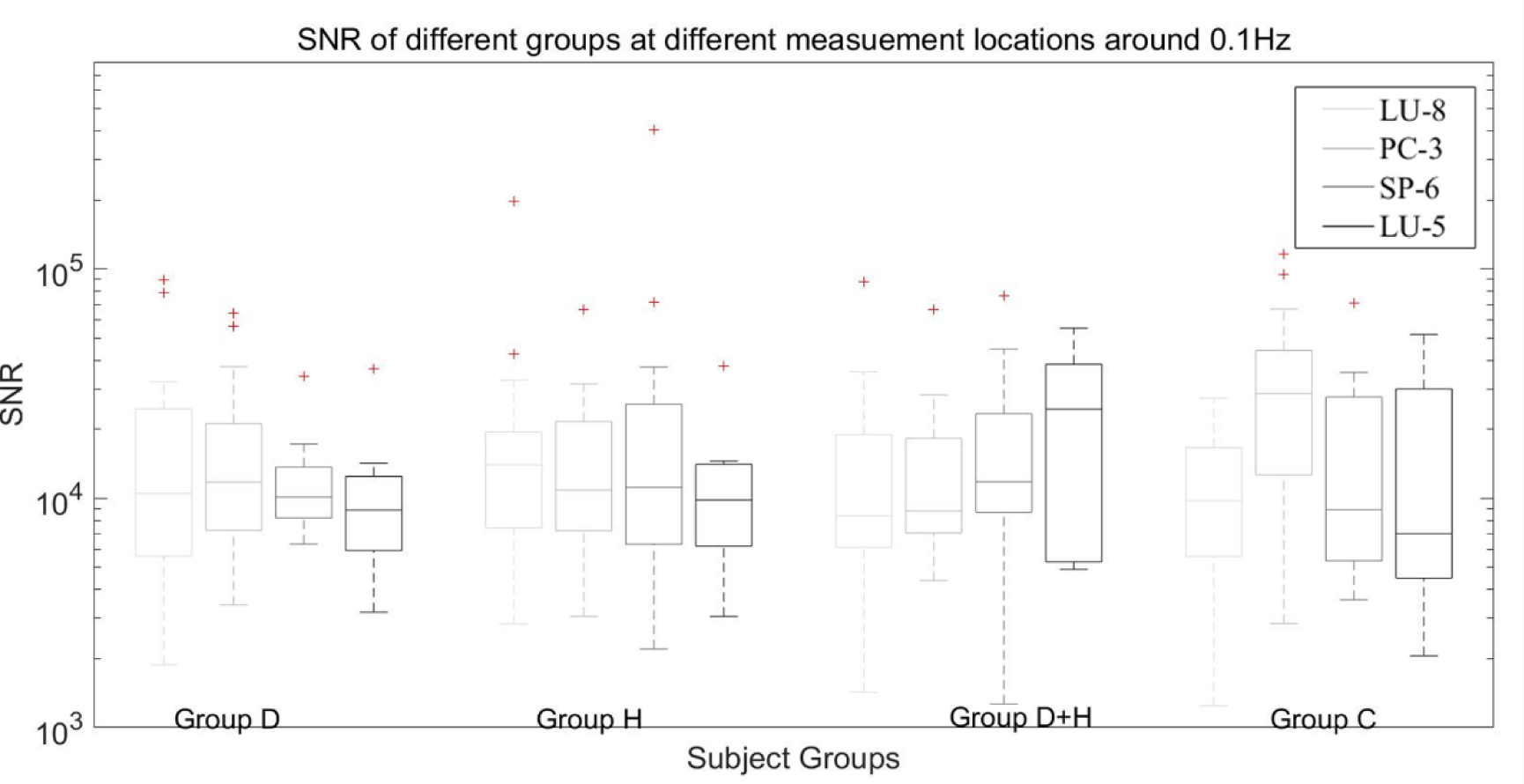
The SNR results of frequency domain about 0.3Hz.

According to Table 4, at the SP-6 location, there is a significant difference between Group H (hypertension) and Group C (control), with a p-value of .0103, which is less than 0.05. Additionally, the difference between Group D (diabetes) and Group C (control) is significant, with a p-value of .0847, which is less than .10. The difference between Group D+H (diabetes + hypertension) and Group C (control) is also significant, with a p-value of .0682, which is less than .10. These results indicate that both diabetes and hypertension significantly influence the SNR of flow motion signals at the SP-6 location in the frequency domain around 0.3Hz.

**Table 4.** SNR of different groups at different measurement locations for frequency band 0.3Hz.

| **p-value** | **LU-8** | **PC-3** | **SP-6** | **LU-5** |
| --- | --- | --- | --- | --- |
| **D/H** | 0.669 | 0.172 | 0.563 | 0.891 |
| **D/D+H** | 0.383 | 0.378 | 0.919 | 0.285 |
| **D/C** | 0.747 | 0.438 | 0.0847 | 0.772 |
| **H/D+H** | 0.413 | 0.758 | 0.446 | 0.348 |
| **H/C** | 0.469 | 0.204 | **0.0103** | 0.889 |
| **D+H/C** | 0.283 | 0.431 | 0.0682 | 0.414 |

Figure 8 displays the SNR values for different groups and measurement locations at a frequency band of 0.1Hz. It illustrates the heterogeneity of SNR at different locations for various diseases at this frequency band.

**Figure 8.**
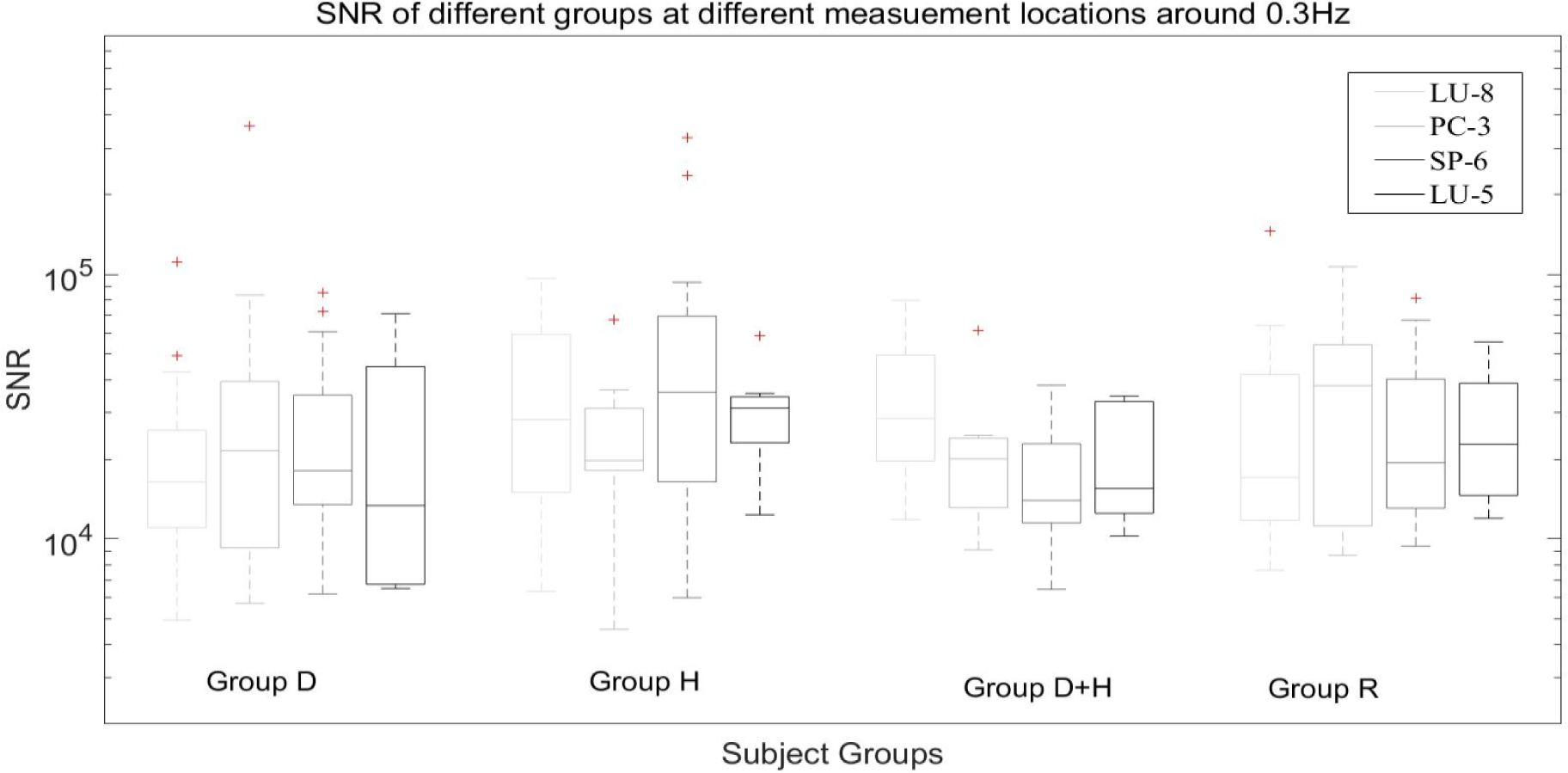
The SNR results of frequency domain about 0.1Hz.

According to Table 5, there is no significant diversity between groups at a frequency of around 0.1Hz. However, at the LU-5 location, there are differences between Group H (hypertension) and Group D+H (diabetes + hypertension) for the 0.1Hz frequency band, with a p-value of .0689, which is less than .10. This suggests that at the LU-5 location, diabetes may influence the SNR results for flow motion signals in patients with hypertension at a frequency band of around 0.1Hz.

**Table 5.** SNR of different groups at different measurement locations for frequency band 0.1Hz.

| ***p*-value** | **LU-8** | **PC-3** | **SP-6** | **LU-5** |
| --- | --- | --- | --- | --- |
| **D/H** | 0.822 | 0.288 | 0.605 | 0.874 |
| **D/D+H** | 0.241 | 0.541 | 0.208 | 0.205 |
| **D/C** | 0.243 | 0.374 | 0.286 | 0.368 |
| **H/D+H** | 0.256 | 0.356 | 0.458 | 0.0689 |
| **H/C** | 0.256 | 0.406 | 0.522 | 0.359 |
| **D+H/C** | 0.978 | 0.745 | 0.948 | 0.470 |

Figure 9 illustrates the SNR values in different groups and measurement locations at the frequency band 0.03Hz. The heterogeneity of SNR at different location for different diseases at the frequency band 0.03Hz.

**Figure 9.**
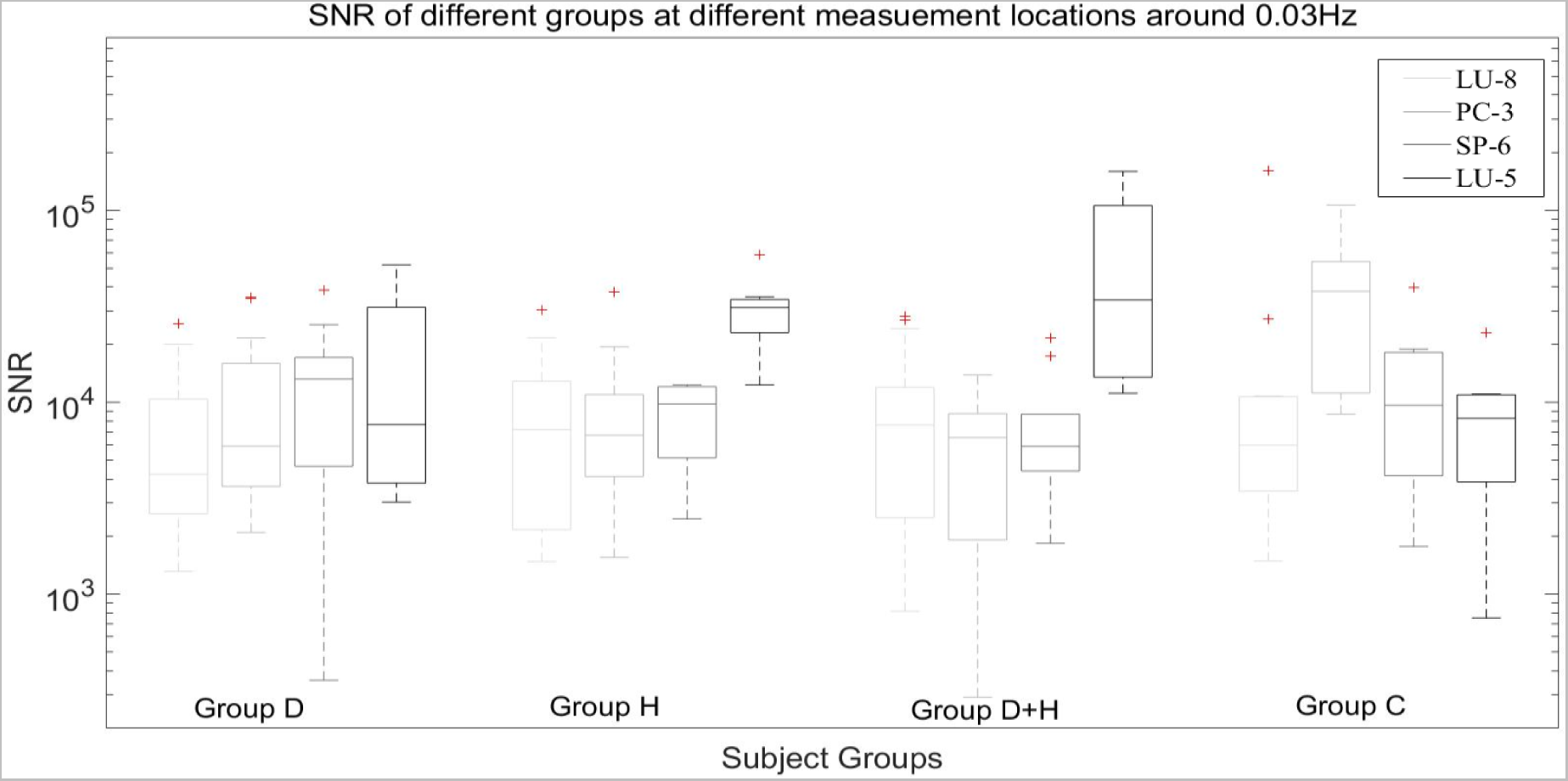
The SNR results of frequency domain about 0.03Hz.

According to Table 6, there is no significant diversity between groups at a frequency of around 0.03Hz. However, at the LU-8 location, there are differences between Group H (hypertension) and Group C (control) for the 0.03Hz frequency band, with a p-value of .0654, which is less than .10. This suggests that at the LU-8 location, hypertension may influence the signal-to-noise ratio (SNR) results for flow motion signals at a frequency band of around 0.03Hz.

**Table 6.** SNR of different groups at different measurement locations for frequency band 0.03Hz.

| ***p*-value** | **LU-8** | **PC-3** | **SP-6** | **LU-5** |
| --- | --- | --- | --- | --- |
| **D/H** | 0.573 | 0.546 | 0.530 | 0.364 |
| **D/D+H** | 0.657 | 0.702 | 0.395 | 0.375 |
| **D/C** | 0.436 | 0.845 | 0.680 | 0.767 |
| **H/D+H** | 0.729 | 0.760 | 0.468 | 0.530 |
| **H/C** | 0.0654 | 0.611 | 0.567 | 0.447 |
| **D+H/C** | 0.168 | 0.795 | 0.158 | 0.410 |

In general, from the noise type results, diabetes may influence the noise color at LU-8, hypertension may influence the noise color of flow motion at LU-8 and PC-3, and the combination of hypertension and diabetes may impact flow motion at LU-8 and SP-6. In this case, LU-8 may reflect the effects of diabetes and/or hypertension on the noise color of flow motion. For the influence of breath activity on vasomotion, the combination of diabetes and hypertension has a significant impact on SNR results at SP-6. For the influence of myogenic activity on vasomotion, diabetes may influence flow motion SNR at LU-5. For endothelial metabolic activity, hypertension may impact SNR of flow motion at LU-8. In summary, vasomotion exhibits different characteristics at different acupoints for individuals in good health as well as those with diabetes or hypertension.LU-5 and LU-8 are both located on the same meridian vessel, the lung meridian of hand-taiyin. However, the manifestation of vasomotion at these two acupoints is not the same. Similarly, PC-3 and LU-5 are located in close proximity to each other, yet the expression of vasomotion at these two acupoints is different. This suggests that the location of an acupoint may not necessarily be related to its expression.

## Conclusion

In summary, this project conducted experimental analysis to determine the effects of diabetes and hypertension on vasomotion. Data was measured using LDF and analyzed using FFT, HHT, and wavelet analysis.

For the LU-8 acupoint, both diabetes and hypertension impact the noise type results, while hypertension impacts the SNR results for endothelial metabolic activity. For the PC-3 acupoint, hypertension may influence the noise type results. For the SP-6 acupoint, the combination of hypertension and diabetes impacts both the noise type results and SNR results for breath. For the LU-5 acupoint, only diabetes impacts the SNR results for myogenic activity.

By comparing the differential results between different measurement groups at different measurement locations, it was found that acupoints have different reflections for both healthy individuals and those with diabetes or hypertension. Regardless of whether acupoints are located on the same meridian vessels or in close proximity to each other, their expression of vasomotion is different.

## Acknowledgments

This work was supported by the Hong Kong Polytechnic University.

## Nonstandard Abbreviations and Acronyms

SNR: Signal Noise Ratio
Ca^2+^: Calcium
NO: Nitric oxide
FFT: Fast Fourier Transformation
SR: Stochastic Resonance
LDF: Laser Doppler Flowmetry
BMI: Body Mass Index
HHT: Hilbert-Huang Transformation
IMFs: Intrinsic Mode Functions

